# Spatial transcriptomics AI agent charts hPSC-pancreas maturation *in vivo*

**DOI:** 10.1101/2025.04.01.646731

**Authors:** Zuwan Lin, Wenbo Wang, Arnau Marin-Llobet, Qiang Li, Samuel D. Pollock, Xin Sui, Almir Aljovic, Jaeyong Lee, Jongmin Baek, Ningyue Liang, Xinhe Zhang, Connie Kangni Wang, Jiahao Huang, Mai Liu, Zihan Gao, Hao Sheng, Jin Du, Stephen J. Lee, Brandon Wang, Yichun He, Jie Ding, Xiao Wang, Juan R. Alvarez-Dominguez, Jia Liu

## Abstract

Spatial transcriptomics has revolutionized our understanding of tissue organization by simultaneously capturing gene expression and spatial localization within intact tissues. However, analyzing these increasingly complex datasets requires specialized expertise across computational biology, statistics, and biological context. To address this challenge, we introduce the Spatial Transcriptomics AI Agent (STAgent), an autonomous multimodal agentic AI that integrates multimodal large language models (LLMs) with specialized computational tools to transform weeks-long analysis tasks into minutes of automated processing. Unlike conventional machine learning approaches that are limited to narrow, predefined tasks, STAgent leverages the emergent capabilities of multimodal LLMs – such as flexible reasoning, contextual understanding, and cross-modal integration – which allow it to adapt to novel data, execute multi-step analyses, and generate biologically meaningful insights with minimal human input. STAgent enables autonomous deep research through integrated capabilities, including dynamic code generation for complex analytical workflows, visual reasoning for interpreting spatial patterns, real-time retrieval of relevant peer-reviewd scientific literature, and synthesis of comprehensive, actionable reports. We applied STAgent to investigate the *in vivo* maturation of human stem cell-derived pancreatic cells (SC-pancreas) transplanted into immunodeficient mice. We generated single-cell spatial transcriptomics data spanning multiple developmental timepoints. STAgent autonomously (1) identified the maturation of initially scattered endocrine cells into well-defined islet-like structures, with predominantly peripheral α-cells surrounding β-cell cores supported by an expanding mesenchymal network; (2) revealed strengthening endocrine-endocrine cell interactions over time and, through context-aware gene set analysis, uncovered spatially resolved biological processes driving maturation; (3) unlike traditional analytical approaches, STAgent offers mechanistic explanations of spatial patterns, contextualizing findings with relevant literatures and developing cohesive insights into human pancreatic development. This agentic approach establishes a new paradigm in spatial transcriptomics analysis by substantially lowering the expertise barrier and reducing analysis time, accelerating biological and biomedical discovery.

## Introduction

Recent advances in spatial transcriptomics technologies have revolutionized the study of tissue organization by enabling simultaneous measurement of gene expressions and their spatial localization within intact tissue sections, providing invaluable insights into complex biological processes such as development, disease progression, and tissue regeneration^1–3^. However, analyzing spatial transcriptomics data remains highly challenging. The high-dimensional nature of these datasets – combining gene expression profiles with spatial coordinates across multiple tissue sections and timepoints – introduces substantial computational complexity that often exceeds the capabilities of conventional analytical approaches.

In addition, interpreting such data typically requires specialized expertise spanning computational biology, statistics, and the relevant biological domain, restricting the broader adoption of these technologies. Current analytical workflows for spatial transcriptomics are often fragmented, time-consuming, and heavily dependent on expert manual intervention, including tool selection, scripting, visualization interpretation, and biological integration. These processes typically take weeks to months per dataset and suffer from limited reproducibility and scalability. As spatial transcriptomics datasets become increasingly complex, analytical bottlenecks have emerged as a major barrier to translating raw data into actionable biological insights.

While traditional machine learning (ML) methods have improved performance in specific tasks such as clustering or cell type and tissue region classification, they lack the core capabilities, including flexible reasoning, contextual understanding, strategic planning, and broad generalization, which are essential for analyzing spatially resolved, multimodal biological data through iterative and multi-step workflows.

Multimodal large language model (LLM) – based AI agents, capable of integrating language, vision, audio, and other data types, offer a promising solution to these challenges. Unlike conventional ML methods, these AI agents exhibit powerful emergent capabilities^4–9^, including advanced reasoning abilities^10,11^, contextual interpretation, and strong potential for zero-shot^12^ or few-shot generalization^9^, enabling them to adapt to novel data and analytical tasks with minimal supervision. Furthermore, their multimodal architecture allows them to reason across diverse data modalities, such as text, code^13,14^, and images^15–17^, in an integrated manner^18,19^, supporting functions such as interpreting visual tissue patterns alongside gene expression data and relevant literature.

Previous reports have demonstrated these capabilities in applications across various scientific disciplines including neural computation^20^, gene editing^21^, and chemistry^22^. Importantly, the ability of such AI agents to reason and synthesize across modalities represents a key distinction from earlier computational approaches and offers significant potential for achieving a more holistic, biologically informed analysis, positioning these agents as powerful tools potentially supporting scientific inquiry and exploration^22–24^. We envision that these integrated capabilities hold the potential to democratize the analysis and interpretation of complex spatial transcriptomics datasets in a scalable manner, reducing dependence on domain expert intervention for each analytical step.

Here, we introduce the Spatial Transcriptomics AI Agent (STAgent), an autonomous multimodal AI agent designed specifically to streamline spatial transcriptomics analysis. STAgent integrates multimodal LLMs with specialized computational tools to perform fully automated, end-to-end analyses without user intervention. The agent’s key capabilities include (1) autonomous code generation for implementing complex analytical workflows; (2) visual reasoning for directly interpreting spatial patterns in tissue architectures; (3) context-aware biological interpretation through real-time scientific literature integration; and (4) comprehensive deep research report generation that transforms raw data into structured output reports detailing methodologies, key findings, biological implications, and literature-supported interpretations. Together, these capabilities reduce time required for spatial transcriptomics analysis from weeks to minutes, dramatically lowering expertise and time barriers to extracting biologically meaningful insights.

To demonstrate STAgent’s capabilities, we applied it to analyze the *in vivo* maturation of human stem cell-derived pancreatic cells (SC-pancreas), a clinically relevant model for diabetes treatment. Diabetes, primarily caused by loss or dysfunction of insulin-producing β-cells, affects hundreds of millions worldwide. Although human pluripotent stem cell (hPSC)-derived pancreatic tissues hold promise for cell replacement therapies, *in vitro*-produced SC-pancreas generally lacks full transcriptional and functional maturity^25,26^. Transplantation into physiologically supportive environments, such as under the kidney capsule, promotes further differentiation and maturation *in vivo*, highlighting key cellular maturation targets that may inform future therapeutic strategies for restoring function lost in diabetes. However, previous studies using single-cell RNA sequencing (scRNA-seq) to analyze these transplanted tissues lacked spatial resolution, leaving important cell composition and cell-cell interactions unresolved^27,28^.

To address this gap, we refined STARmap^29^, an imaging-based spatial transcriptomics approach, to profile SC-pancreas grafts across multiple maturation stages in immunodeficient mice. STAgent autonomously analyzed this high-dimensional spatiotemporal gene expression dataset, uncovering the progressive organization of endocrine cells into well-structured islet-like clustered characterized by mantle-core α-β cell patterns supported by an expanding mesenchymal network. The agent also detected enhanced endocrine cell interactions over time, indicative of islet spatial maturation. In addition, STAgent’s context-aware gene set analysis further identified key biological processes driving maturation, including enhanced ion transport, secretory granule formation, and cell-matrix interactions.

This AI agent-driven approach offers a scalable paradigm for spatial transcriptomics analysis applicable across living systems and processes. By automating complex, multi-step analytical tasks and integrating biological context, STAgent substantially reduces the need for specialized expertise and accelerates the transition of spatial biological data into scientific discoveries, offering a scalable and generalizable platform across diverse biomedical research domains.

## Results

### Spatial transcriptomics of SC-pancreas grafts and AI agent development

Spatial transcriptomics in pancreatic tissue has historically been limited by technical challenges related to tissue processing and signal sensitivity. To address this, we developed a modified STARmap protocol specifically optimized for pancreatic tissue, building upon methods described by Farack *et al*.^28^. Key optimizations include extending mRNA denaturation step to at least 3 hours and increasing the formamide concentration from 10% to 30% during hybridization (**Extended Data** Fig. 1a). These optimizations enabled sensitive, single-molecule RNA detection in pancreatic tissues comparable to STARmap performance in other tissue types.

Using established protocols^25–27^, we generated human SC-pancreas cells and transplanted them under the kidney capsule of immunodeficient mice. We collected grafts at 4-, 16-, and 20-week post-transplantation to capture distinct stages of *in vivo* maturation (**Fig. 1a**). Using our optimized STARmap protocol (**Fig. 1a** and Extended Data Fig. 1a, b), we profiled over 60,000 cells across tissue sections spanning all three timepoints. Multi-round *in situ* sequencing data were computationally registered and segmented into 3D-resolved single cells, producing a high-fidelity, spatially preserved cell-by-gene expression matrix for downstream analysis (**Fig. 1b**).

**Fig. 1:**
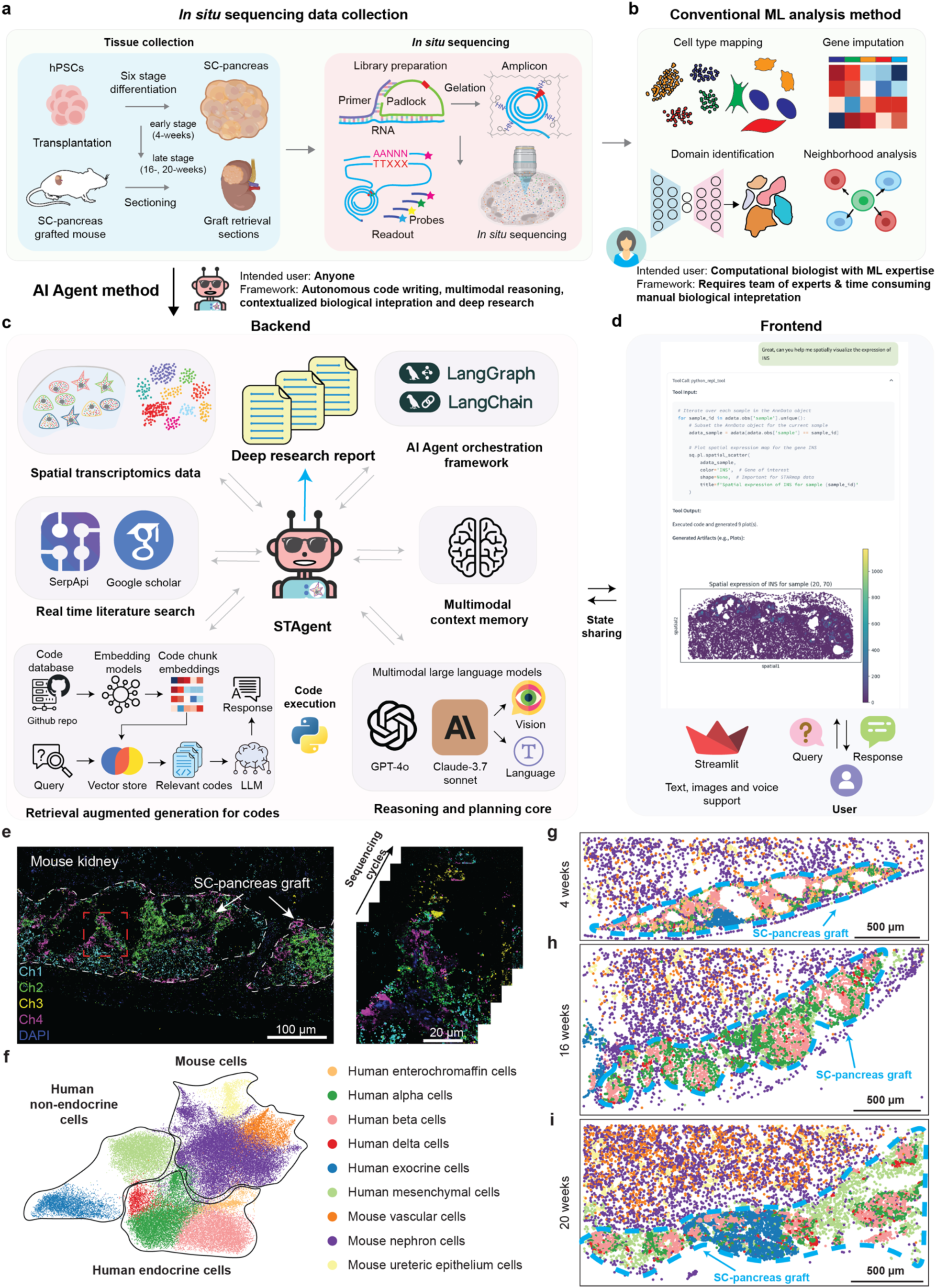
Spatial transcriptomics workflow and multimodal AI agent architecture. (**a**) Experimental workflow for SC-pancreas graft analysis. hPSCs are differentiated into SC-pancreas and transplanted under the kidney capsule of immunodeficient mice, with grafts collected at early (4 weeks) and late (16-20 weeks) stages. *In situ* sequencing includes padlock probe hybridization, rolling circle amplification, and multi-round fluorescent sequencing for spatial gene expression detection. (**b**) Conventional machine learning-based analysis for spatial transcriptomics requires substantial computational expertise and manual interpretation. (**c**) STAgent backend architecture integrates spatial transcriptomics data, real-time literature search, retrieval-augmented code generation, multimodal large language models for reasoning and planning, multimodal context memory, and an AI agent orchestration framework to generate comprehensive research reports. (**d**) STAgent frontend provides an intuitive user interface for multimodal interaction, delivering rich visual and analytical outputs. (**e**) Representative raw fluorescence images from *in situ* sequencing experiments showing engrafted SC-pancreas. (**f**) UMAP visualization illustrating cell clustering results, highlighting distinct cell populations, including human endocrine (α-, β-, δ-, enterochromaffin), human non-endocrine (exocrine, mesenchymal), and mouse (vascular, nephron, ureteric epithelium). (**g**–**i**) Spatial transcriptomics maps at 4-(**g**), 16-(**h**), and 20-weeks (**i**) post-transplantation. Each point represents an individual cell, color-coded by major cell type following the scheme defined in panel (**f**).

Despite advances in spatial transcriptomics technologies, the interpretation of such high-dimensional spatiotemporal data remains a major bottleneck. Current workflows require extensive expert involvement across multiple stages, including analytical tool selection, parameter running, result interpretation, and integration data across multiple timepoints and tissue slices. This requirement makes the process not only time-consuming, but also subjective and difficult to scale. These are particularly limited in analyzing dynamic systems such as pancreatic islet maturation, where rapid and complex changes in tissue architecture, cell composition and cell-cell interactions demand both precise and flexible analysis. Therefore, there is a critical need for an autonomous, scalable analytical framework capable of generating consistent and biologically meaningful insights across diverse and evolving datasets.

To address this challenge, we developed STAgent, an autonomous multimodal AI agent designed for spatial transcriptomics analysis (**Fig. 1c-d**). STAgent comprises a backend reasoning engine and an intuitive user-facing frontend, connected via an orchestration layer that manages data flow and analytical operations.

The backend (**Fig. 1c**) integrates (1) multimodal LLMs (Claude-3.7-Sonnet^30^, GPT-4o^31^) capable of processing both textual and visual inputs; (2) a multimodal context memory system that tracks analysis history and maintaining consistency; (3) a retrieval-augmented code generation (RAG)^32^ module that dynamically generates analytic scripts from a Chroma-indexed GitHub codebase^33^; and (4) a real-time literature integration module that contextualizes STAgent’s findings in peer-reviewed scientific knowledge.

A defining feature of STAgent is its visual reasoning capability. Unlike conventional tools that operate on numerical matrices, STAgent can directly interpret spatial transcriptomic images and cell type maps. This allows STAgent to autonomously identify tissue architectures, spatial arrangement of cell types, and microenvironmental patterns – insights that typically require extensive expert review. STAgent can also detect spatial relationships, developmental transitions, and cell-cell interactions that are difficult to quantify using traditional methods. Moreover, by using deep contextual knowledge, STAgent can detect subtle patterns and generate biologically plausible hypotheses, which cannot be generated from traditional manual analysis alone.

To support its RAG capabilities, we implemented a pipeline that scrapes public code repositories, segments them into analyzable units, and embeds them into a vector database. When presented with a novel analysis task, STAgent queries this database to retrieve and refine relevant code templates using the LLM, thus enabling the performance of complex analytical workflows without manual programming.

The frontend interface (**Fig. 1d** and Extended Data Fig. 2), built in Streamlit^34^, allows users to interact with STAgent using natural language, images, or even voice commands. Orchestration of these components is managed through LangGraph and LangChain^35^, ensures smooth coordination between components and adaptive behavior across datasets.

STAgent’s most transformative feature is its ability to autonomously generate structured research reports (**Fig. 1c**). These reports are organized like traditional scientific publications, including sections on methodology, key findings, biological interpretations, and literature-supported discussion, but generated entirely without human intervention. By integrating real-time literature search tools (e.g., SerpAPI, Google Scholar)^36^ (**Fig. 1c**), STAgent ensures that all interpretations are transparently grounded in peer-reviewed scientific knowledge. This drastically accelerates the translation of complex spatial transcriptomics datasets into coherent, biologically meaningful insights, compressing what traditionally required weeks of expert manual analysis into minutes.

### Conventional machine-learning-based spatial analysis of graft development

To establish a robust interpretive baseline for evaluating STAgent’s AI-driven workflow, we first performed conventional ML–based analyses on spatial transcriptomocs data following five cycles of *in situ* sequencing (**Fig. 1e**). Initial clustering identified major human pancreatic cell types – endocrine (α, β, δ and enterochromaffin cells) and non-endocrine (exocrine, mesenchymal) – alongside mouse host kidney cells (vascular, nephron, ureteric epithelium) (**Fig. 1f**). Mapping these cell types to spatial coordinates generated comprehensive cell-type maps, illustrating the evolving graft architecture over time (**Fig. 1g-i**). At 4 weeks post-transplantation, the graft cells formed scattered clusters (**Fig. 1g**). By 16 weeks, cells began assembling into organized structures, including endocrine cells aggregating into early islet-like formations (**Fig. 1h**). By 20 weeks, grafts exhibited clearly defined islet-like structures, characterized by intermingled α-and β-cells surrounded by expanding mesenchymal populations (**Fig. 1i**).

Each cell type displayed distinct gene expression signatures (**Extended Data** Fig. 3): β-cells expressed insulin (INS); α-cells, glucagon (GCG); δ-cells, somatostatin (SST); and enterochromaffin cells, TPH1. Exocrine and mesenchymal cells specifically expressed CPA2/CTRB2 and COL1A1/COL3A1, respectively. Mouse kidney host cells exhibited tissue-specific markers SPP2/LRP2 (nephron), AQP2 (ureteric epithelium), and CDH5/KDR (vascular).

To characterize complex spatial patterns beyond single-cell identities, we applied STAligner^37^, a graph attention autoencoder integrating spatial and gene expression data across graft slices (**Extended Data** Fig. 4a). This analysis defined six distinct spatial domains: four primarily human-derived (h1-h4) and two predominantly mouse-derived (m1-m2) (**Extended Data** Fig. 4b-e). Each human domain had a distinct cellular composition (**Extended Data** Fig. 5a-c): α-cell–enriched endocrine (h1), β-cell–enriched endocrine (h2), exocrine-like (h3), and mesenchymal (h4). Early grafts showed intermingled domains, indicative of immature cellular expansion and active remodeling, while maturation resulted in increasingly segregated domains with compact endocrine clusters resembling native islet architecture.

Quantitative assessment confirmed these developmental patterns (**Extended Data** Fig. 5d). At 4 weeks, α-cell (h1) and β-cell (h2) domains had fewer endocrine cells than expected, whereas the mesenchymal domain (h4) was already evident. By 16-20 weeks, substantial α- and β-cell expansion indicated greater commitment to endocrine lineage development, accompanied by prominent mesenchymal growth.

Spatial neighborhood analyses revealed progressive emergence of cell-type-specific interactions (**Extended Data** Fig. 5e). At 4 weeks, cell-cell interactions appeared diffuse with less preferential associations. By 16 weeks, interactions among endocrine cells strengthened, becoming particularly prominent by 20 weeks. At this stage, positive enrichment for α-α, β-β, and α-β cell interactions indicated functional maturation of islet-like structures capable of coordinated glucose regulation.

To expand analyses beyond the original 154-gene panel, we used Tangram^38^ for genome-wide spatial expression imputation based on integration with a reference scRNA-seq atlas, increasing coverage to over 15,000 genes (**Extended Data** Fig. 6a). Validation confirmed accurate recapitulation of original STARmap gene expression patterns (**Extended Data** Fig. 6b). The expanded dataset revealed spatially resolved expression of key pancreatic development genes absent from the original panel, such as INS-IGF2 (β-cell development), PCSK2 (proinsulin processing), PRSS1 (exocrine function), and COL1A2 (extracellular matrix formation).

Finally, we characterized intercellular signaling evolution using CellChat^39^ on the imputed dataset. Signaling network connectivity markedly increased from 4 to 20 weeks (**Extended Data** Fig. 6c, d), reflecting enhanced functional maturity necessary for islet coordination. Specifically, endocrine cell-cell communication intensified over time (**Extended Data** Fig. 6e), aligning consistent with the formation of tightly interconnected islet-like structure. Notably, endocrine-mesenchymal interactions also strengthened, underscoring potential stromal support for islet maturation.

### STAgent enables autonomous spatial transcriptomics analysis

Although the conventional ML-based spatial transcriptomics analyses described above provided valuable insights into SC-pancreas graft development. Their implementation poses significant practical challenges: they typically require extensive cross-disciplinary expertise and manual coding skills, with the interpretive analysis of spatial data representing the most crucial bottleneck. Domain experts must integrate gene expression patterns, spatial coordinates, and cellular interactions across multiple tissue sections – a complex, subjective, and time-intensive task that can extend over weeks to months. This expertise barrier severely limits the scalability and accessibility of spatial transcriptomics studies, creating an urgent need for automated analytical tools.

To address this, we test the feasibility and the reasoning capabilities of STAgent to perform spatial transcriptomics analysis without human intervention. By leveraging the emergent capabilities^7^ of the agent, particularly its dynamic reasoning and code generation^13^, we evaluated whether STAgent could autonomously navigate the multi-step workflow that traditionally demands domain expertise. Given our pancreatic islet development dataset and a straightforward natural language instruction to “*perform an end-to-end analysis*,” STAgent executed a comprehensive analytical workflow entirely autonomously (**Fig. 2**). The agent automatically generated Python code tailored to each analytical step (**Fig. 2a**), employing specialized bioinformatics libraries, including *Scanpy*^40^ and *Squidpy*^41^, for sophisticated analyses.

**Fig. 2:**
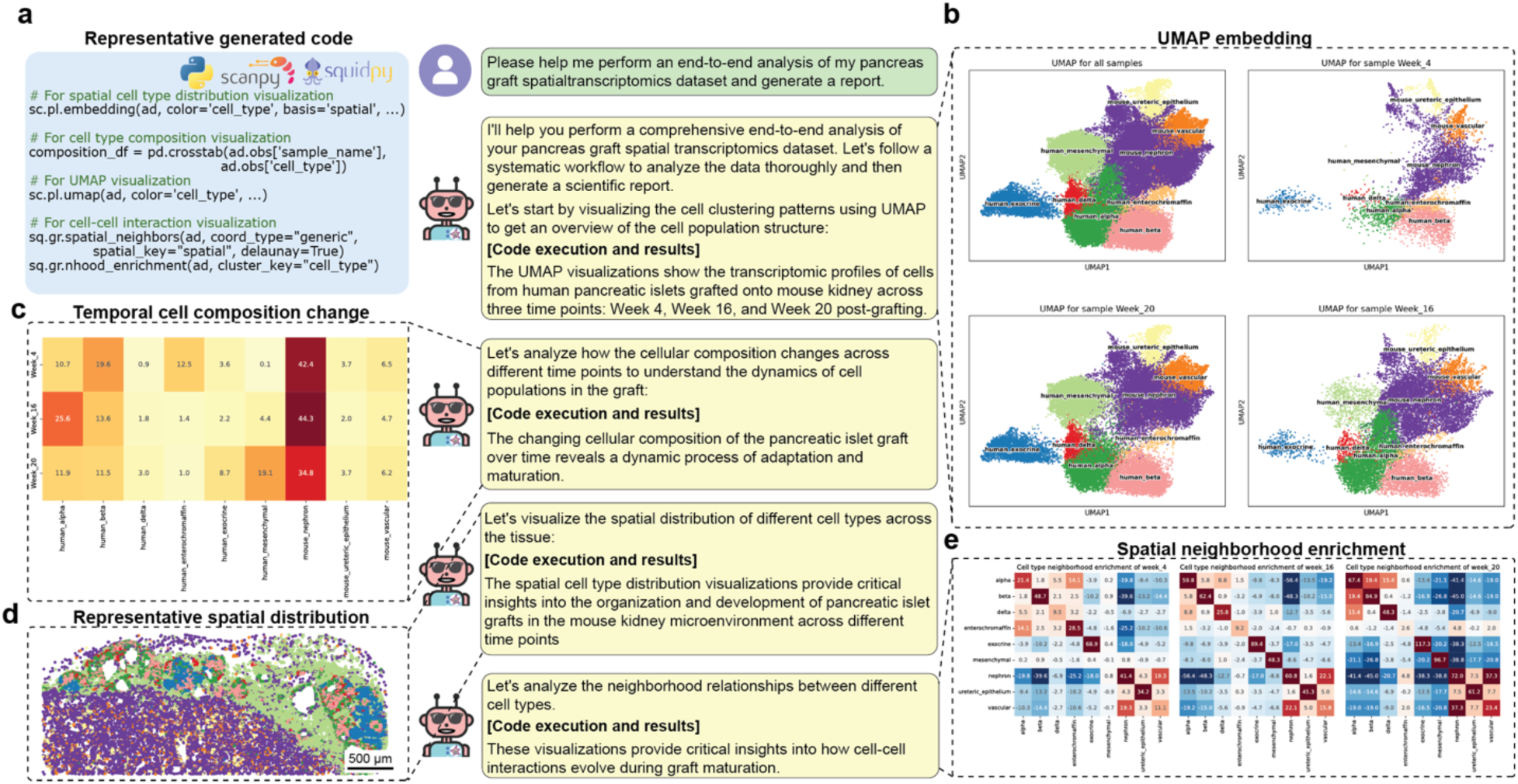
STAgent enables autonomous analytical processing of spatial transcriptomics data. (**a**) STAgent autonomously generates Python code for spatial transcriptomics analysis using specialized libraries in response to natural language requests, eliminating the need for manual programming. (**b**) STAgent autonomously generates UMAP embeddings for visualizing transcriptional profiles across three developmental timepoints, revealing distinct cell populations and their relationships. (**c**) STAgent autonomously generates temporal cell composition analysis showing changing proportions of cell types during graft maturation. (**d**) STAgent autonomously generates spatial maps of cell-type distributions, providing insights into the evolving spatial architecture of human SC-pancreas grafts within the mouse kidney microenvironment. (**e**) STAgent autonomously performs spatial neighborhood enrichment analysis revealing temporal evolution of cell-cell interactions. Heatmaps depict preferential associations among cell types across different developmental stages.

STAgent’s autonomous analytical pipeline proceeded in a structured manner. First, it generated UMAP embeddings to visualize global transcriptional profiles across developmental timepoints, offering an immediate overview of evolving cell populations (**Fig. 2b**). Next, the agent quantified temporal changes in cellular composition, creating heatmaps that clearly captured dynamic shifts in cell-type proportions as grafts matured (**Fig. 2c**). STAgent then mapped identified cell types onto spatial coordinates, producing comprehensive distribution maps illustrating the architectural development of pancreatic grafts within the mouse kidney microenvironment over time (**Fig. 2d**). Lastly, STAgent analyzed spatial neighborhood relationships among cell types, generating spatial enrichment heatmaps to elucidate evolving cell-cell interactions during graft maturation (**Fig. 2e**). Taken together, these results underscore how emergent multimodal AI agents can streamline otherwise labor-intensive workflows and deliver rapid, expert-level analysis into complex spatial transcriptomics data.

### STAgent enables visual reasoning and contextualized biological interpretation

Beyond automating technical workflows, we next evaluated whether STAgent’s visual reasoning abilities could autonomously identify subtle biological features from spatial transcriptomics data, an interpretive task generally requiring expert knowledge. In contrast to conventional tools restricted to numerical inputs, STAgent can integrate vision-language reasoning^15–17^ to directly interpret spatial visualizations, thereby detecting morphological patterns and tissue architectures that often go unnoticed without extensive manual inspection (**Fig. 3**). By transforming these traditionally subjective, time-consuming interpretive tasks into objective, scalable, and automated processes, STAgent could fundamentally redefine how spatial transcriptomics can be explored at scale.

**Fig. 3:**
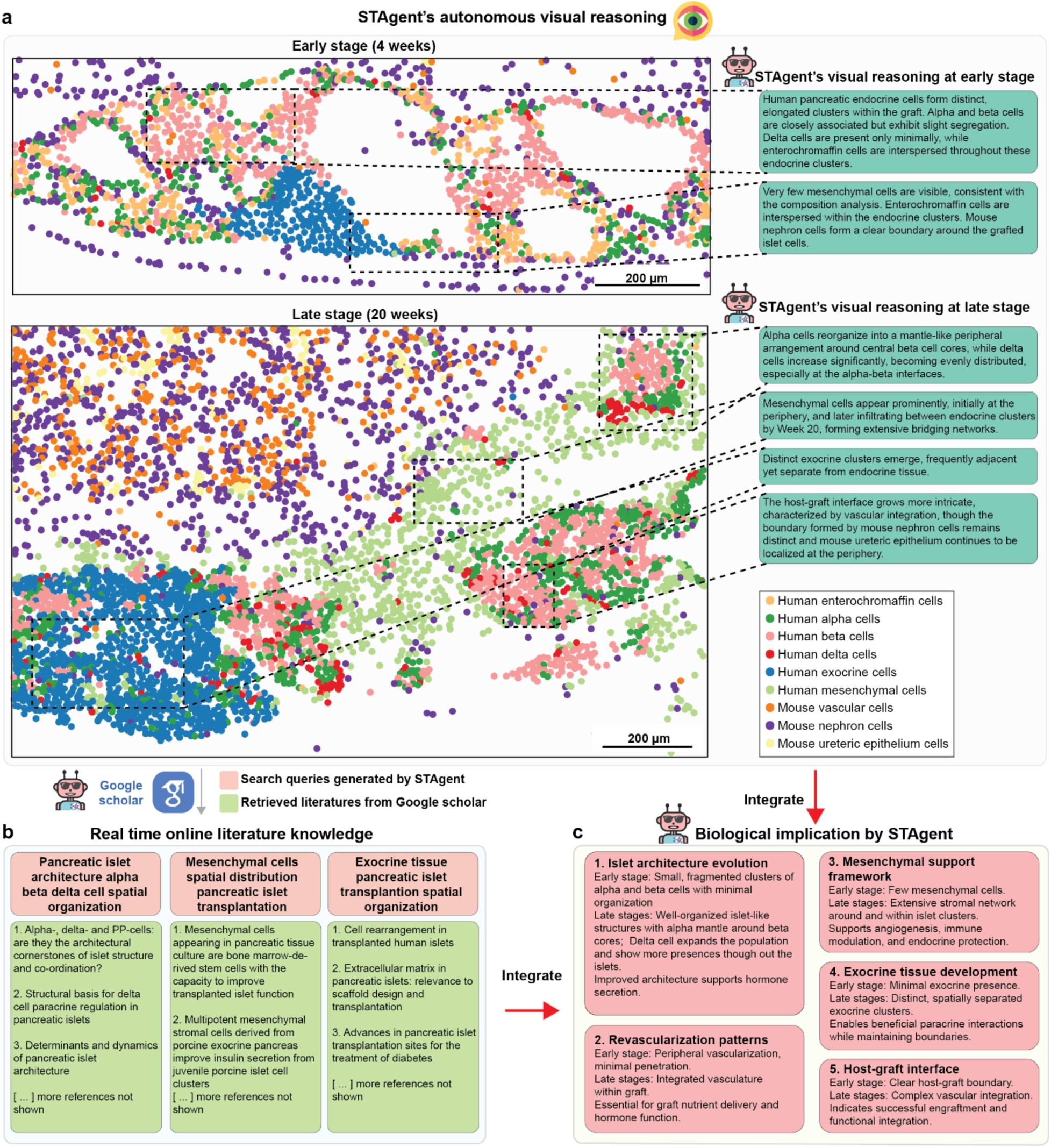
STAgent enables autonomous visual reasoning and contextualized biological interpretation. (**a**) Autonomous visual reasoning performed by STAgent, analyzing spatial distributions of cell types at early (4 weeks) and late (20 weeks) graft development stages. At the early stage, STAgent identifies elongated endocrine cell clusters with minimal segregation between α- and β-cells, few δ-cells, and a distinct boundary formed by mouse nephron cells. At the late stage, STAgent detects reorganization of α-cells into a mantle-like arrangement around β-cell cores, increased δ-cells at α-β interfaces, prominent mesenchymal cell networks, distinct exocrine clusters, and more intricate graft-host vascular integration. (**b**) Autonomous literature search capability, exemplified by STAgent-generated queries and retrieval of relevant scientific literature from Google Scholar, contextualizing visual analyses. (**c**) Synthesis of biological implications through integration of visual reasoning with literature-based knowledge. STAgent identifies early and late-stage developmental patterns and infers functional significance.

At the early graft stage (4 weeks), STAgent visually identified elongated clusters of human endocrine cells, noting α- and β-cells closely associated yet slightly segregated (**Fig. 3a, top**). The agent highlighted the minimal presence of δ-cells and scattered enterochromaffin cells, along with the limited distribution of mesenchymal cells and a clear graft-host boundary delineated by mouse nephron cells. At the late stage (20 weeks), STAgent recognized substantial developmental changes (**Fig. 3a, bottom**): α-cells reorganized into peripheral mantle-like arrangements surrounding central β-cell cores; δ-cells increased significantly, becoming evenly distributed, particularly at α-β interfaces; mesenchymal cells emerged prominently, infiltrating endocrine clusters from the periphery and forming extensive stromal networks; distinct exocrine clusters developed adjacent endocrine tissue; and the graft-host interface became increasingly complex through integrated vascularization.

Importantly, STAgent integrates its visual analysis with automated literature retrieval (**Fig. 3b**), enabling immediate contextualization of results within established knowledge on pancreatic islet biology (**Fig. 3c**) – an interpretive process traditionally requiring extensive expert training. The agent connected visual observations to documented developmental processes, such as the evolution of islet architecture facilitating coordinated hormone secretion, the progressive establishment of vascular networks essential for nutrient delivery, and the formation of mesenchymal frameworks that support angiogenesis, immune modulation, and endocrine stability. Additionally, STAgent underscored the biological relevance of spatially segregated exocrine clusters facilitating beneficial paracrine interactions while preserving tissue compartments. Finally, STAgent emphasized the transformation of the graft-host interface into integrated vasculature indicative of successful functional engraftment, supporting these conclusions with relevant literature identified autonomously.

To further assess STAgent’s biological reasoning depth beyond spatial visualization, we evaluated its performance in context-aware gene set analysis (**Fig. 4**). Traditional enrichment analyses often produce biologically irrelevant pathways, especially when applied to specialized tissues like pancreatic islets (**Fig. 4a, top**) due to their sole reliance on statistical overrepresentation from generic databases. In contrast, STAgent employs context-aware multimodal reasoning informed by targeted literature searches to provide biologically grounded interpretations (**Fig. 4a, bottom**).

**Fig. 4:**
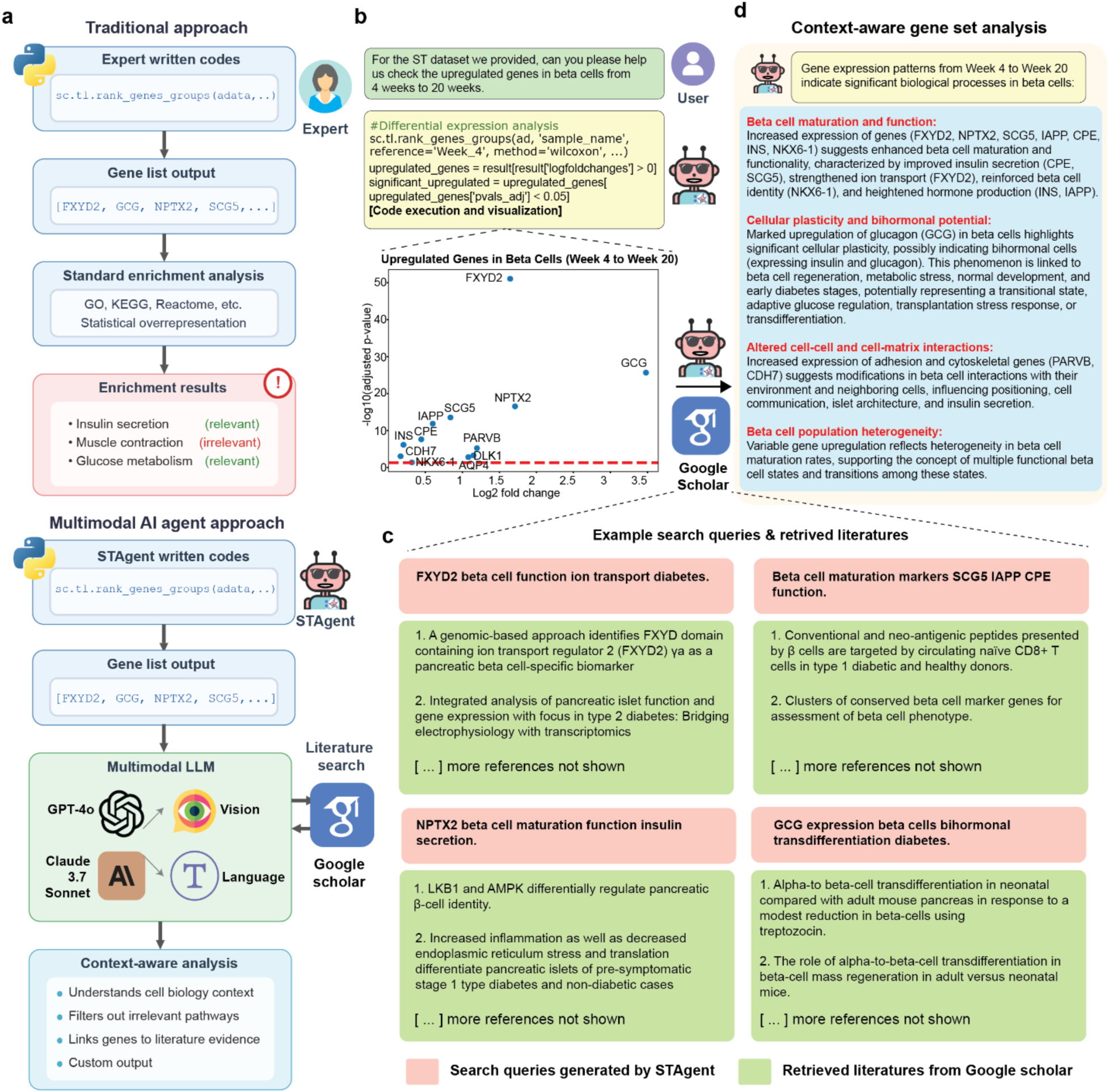
STAgent enables context-aware gene set analysis. (**a**) Comparative schematic contrasting traditional gene-set analysis methods (top) with STAgent’s multimodal, context-aware approach (bottom). Traditional methods require manual expert-driven coding, produce gene lists, and rely on standard enrichment tools that often yield biologically irrelevant processes. STAgent autonomously generates tailored code, interprets gene lists through multimodal LLMs, and produces context-aware analyses specific to pancreatic biology. (**b**) STAgent’s autonomous differential gene expression analysis of β-cells between 4 and 20 weeks visualized as a volcano plot, highlighting key developmental markers. (**c**) Example search queries autonomously generated by STAgent and associated literature retrieved from Google Scholar, contextualizing identified differentially expressed genes in the biology of pancreatic β-cells. (**d**) Autonomous biological interpretation organized by STAgent, categorizing gene expression changes into functional groups with detailed, context-driven insights.

When analyzing differential gene expression between 4- and 20-week β-cells, STAgent autonomously generated and executed custom code to identify differentially expressed genes and visualize their distribution (**Fig. 4b**). Importantly, instead of relying solely on statistical enrichment results, STAgent contextualized these findings within the framework of pancreatic β-cell biology, integrating multimodal reasoning with scientific literature retrieved autonomously from Google Scholar (**Fig. 4c**). STAgent then categorized the results into functionally relevant themes relevant to β-cell maturation (**Fig. 4d**): (1) enhanced β-cell maturation and insulin secretion; (2) cellular plasticity and bihormonal potential indicated by GCG upregulation; (3) altered cell-cell and cell-matrix interactions influencing β-cell microenvironment integration; and (4) emerging β-cell population heterogeneity, reflective of functional diversification during maturation. This targeted, context-aware approach filtered irrelevant pathways typically seen in conventional analyses, producing biologically meaningful insights into β-cell development and maturation.

Collectively, these results confirm that STAgent not only automated the technical components of spatial transcriptomics data analysis but also delivers robust, expert-level biological intrepretations of tissue organization and gene function through advanced visual reasoning and context-aware analytics. By drastically reducing the time and expertise required for such complex analysis, STAgent offers a scalable, intelligent framework for understanding spatial biology.

### STAgent enables autonomous deep research and knowledge synthesis

Building on its capacity for autonomous multimodal analysis and contextualized biological interpretation, we next evaluated whether STAgent could autonomously transform complex spatial transcriptomics data into cohesive, publishable research reports (**Fig. 5**). Unlike conventional analytic pipelines for such tasks require extensive domain expertise and weeks to months of manual efforts, STAgent integrates all the previously demonstrated components into a streamlined workflow that performs autonomous data ingestion, dynamic code generation, visual-language reasoning, targeted literature retrieval, and the final assembly of findings into a coherent scientific report.

**Fig. 5:**
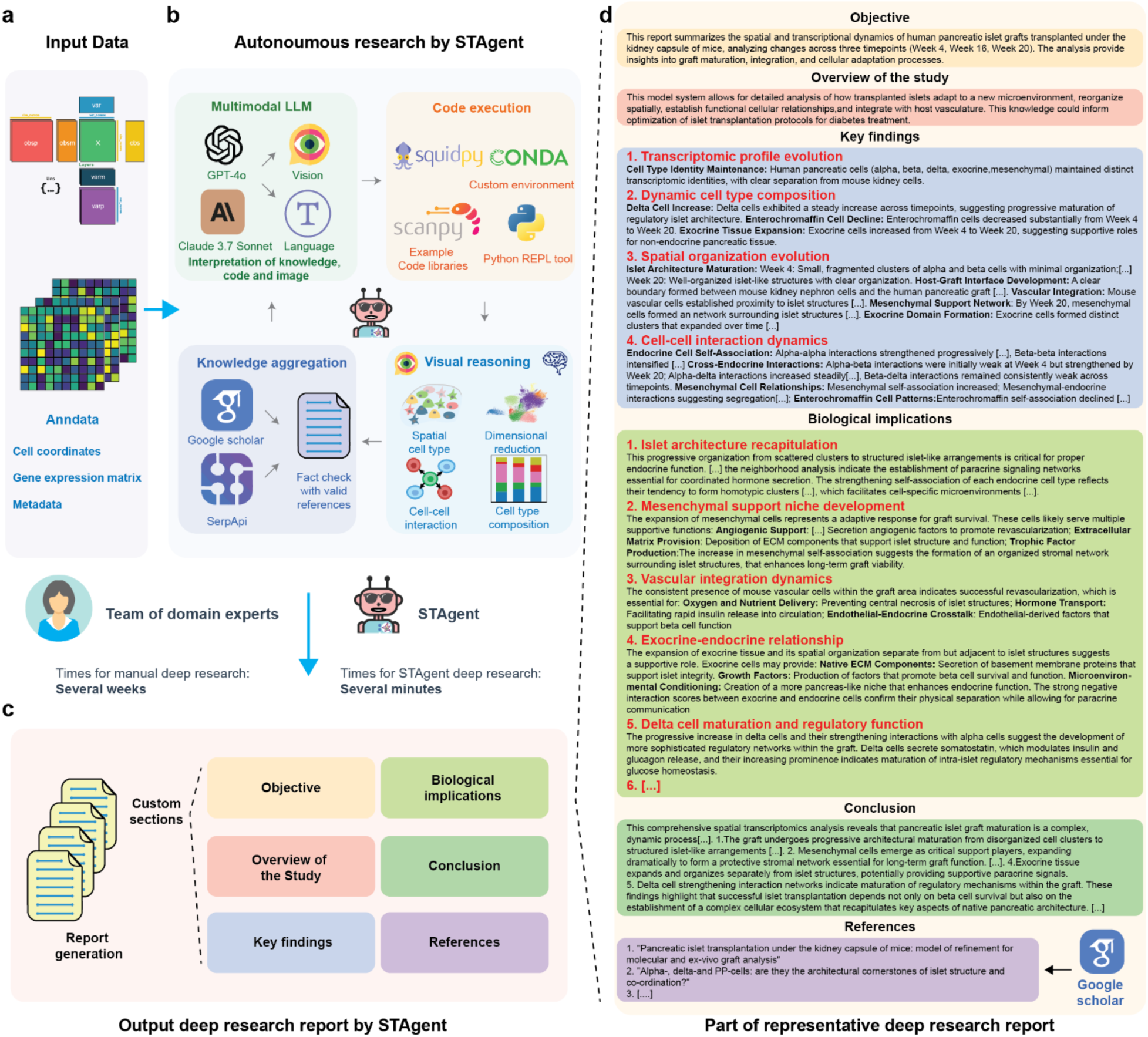
STAgent enables autonomous end-to-end deep research and knowledge synthesis. (**a**) Input spatial transcriptomics data consisting of cell coordinates, gene expression matrices, and metadata. (**b**) Overview of STAgent’s autonomous research pipeline, leveraging multimodal LLMs to interpret data, generate specialized analytical code, aggregate insights from literature, and apply visual reasoning to detect spatial patterns. (**c**) Autonomous generation of comprehensive research reports by STAgent, synthesizing and consolidating knowledge from spatial analyses, literature context, and visual reasoning. Structure of autonomously generated report includes clearly defined sections: Objective, Overview, Key Findings, Biological Implications, Conclusion, and References. (**d**) Representative example of a research report generated autonomously by STAgent, showing its capacity for coherent, detailed, context-rich scientific communication.

When provided only with spatial transcriptomics data (**Fig. 5a**) and a high-level objective – analyzing pancreatic graft maturation – STAgent autonomously developed and executed a complete research workflow. This involved integrating multiple data modalities, including spatial coordinates, gene expression matrices, and metadata (**Fig. 5a**), applied multimodal reasoning to interpret patterns and relationships within the datasets, and retrieved relevant literature from Google Scholar (**Fig. 5b**). This research workflow produced a well-structured research report (**Fig. 5c, d**), organized similarly to conventional scientific publications, including sections for findings into clearly defined sections: *Objective, Overview of the Study, Key Findings, Biological Implications, Conclusion, and References*.

Specifically, in the *Key Findings* section (**Fig. 5d**), STAgent synthesized results from its multimodal analyses to identify developmental patterns such as transcriptomic profile evolution, dynamic changes in cell composition, spatial organization, and evolving cell-cell interaction dynamics. The *Biological Implications* section interpreted the functional significance of these findings for pancreas development, covering islet architecture recapitulation, mesenchymal niche formation, vascular integration, exocrine-endocrine relationships, and intra-islet communication. STAgent’s interpretations extended beyond description to propose mechanisms – for instance, suggesting that progressive structural organization is critical for endocrine function and coordinated hormone secretion, and that increasing mesenchymal self-association indicates stromal network formation crucial for long-term graft viability. Such insights typically require specialized pancreatic development expertise. Throughout the report, STAgent autonomously integrated citations to contextualize findings within broader scientific literature. The *Conclusion* section consolidated these findings into a cohesive narrative, emphasizing interactions among cell types and progressive native-like tissue organization. Three illustrative examples demonstrate STAgent’s ability to autonomously synthesize knowledge and interpret complex data:

First, STAgent analyzed the spatiotemporal progression of endocrine cell organization within pancreatic grafts. At 4 weeks, analyses showed fragmented and disorganized α-, β-, and δ-cell clusters indicative of immature islet architecture. By 16 weeks, STAgent identified significantly improved spatial coherence with larger endocrine cell groupings. By 20 weeks, fully matured islet-like structures with organized α-β-δ arrangements emerged. Quantitative cell-cell interaction analyses validated progressively stronger α-β and α-δ interactions linked to functional maturation. To contextualize these findings, STAgent autonomously retrieved supportive literature (e.g., Arrojo e Drigo et al^42^, Brereton et al^43^), confirming similar patterns where reorganized islet architecture enabled coordinated hormone secretion and concluding that architectural maturation promotes endocrine functionality and promotes long-term graft success.

Second, through UMAP dimensionality reduction and cell composition analyses, STAgent observed a substantial mesenchymal cell expansion, from 4 to 20 weeks. Spatial analyses demonstrated mesenchymal cells progressively formed extensive networks encircling and penetrating endocrine structures, suggesting adaptive stromal responses. Literature retrieved by STAgent (e.g., Hematti et al^44^, Barachini et al^45^, Sordi et al^46^) highlighted mesenchymal cells’ critical roles in revascularization, immunomodulation, and trophic support. Integrating these insights, STAgent concluded mesenchymal cell expansion enhances graft stability by fostering supportive microenvironments necessary for long-term functionality.

Third, STAgent examined host-driven vascularization by analyzing spatial organization evolution and cell-cell interactions. At 4 weeks, vascularization was primarily peripheral, characterized by mouse vascular cells surrounding – but minimally infiltrating – human endocrine clusters. By 16 weeks, STAgent detected active angiogenesis, with mouse vessels increasingly penetrating the graft and found this vascular pattern matured by 20 weeks into an integrated network that infiltrated endocrine structures. Based on the retrieved literature from Pepper et al.^47^, STAgent confirmed a two-stage islet revascularization process: initial peripheral angiogenesis followed by intra-islet vascular integration. The agent interpreted this transition as critical for delivering essential oxygen and nutrients to the developing islets and for constituting a critical pathway for hormone secretion into systemic circulation – a key determinant of transplantation success.

Collectively, these examples demonstrate STAgent’s capability for multi-layered interpretation beyond routine data processing. By synthesizing multimodal data, generating biologically grounded insights, and producing structured scientific outputs – entirely without human input – STAgent represents a transformative approach to spatial transcriptomics analysis. It democratizes access to complex data interpretation and significantly accelerates scientific discovery across diverse research fields.

## Discussion

In this study, we introduced STAgent, an autonomous multimodal AI agent for spatial transcriptomics analysis, and applied it to investigate the *in vivo* maturation of SC-pancreas. By integrating high-resolution spatial transcriptomics with autonomous AI agent-driven analysis, STAgent not only provides deep insights into SC-pancreas maturation but also establishes a powerful and scalable framework for analyzing and exploring complex biological systems.

The foundation of STAgent’s capabilities lies in the emergent properties of multimodal LLMs. Unlike conventional machine learning methods that are often constrained to narrow, task-specific functions, multimodal LLMs exhibit emergent abilities such as dynamic reasoning, multimodal understanding, and context-aware decision-making. These capabilities arise not from explicit programming, but from the scale and diversity of training data, which enable the models to generalize across domains and perform complex, previously unanticipated tasks. By leveraging these emergent features, STAgent transcends traditional analytic pipelines – it not only executes technical workflows but also interprets spatial tissue architectures, contextualizes biological significance, and synthesizes findings into coherent scientific narratives. This shift from algorithmic execution to autonomous reasoning is what enables STAgent to operate as a true scientific collaborator rather than a passive analytical tool. We summarize STAgent’s core capabilities below, highlighting how it transforms spatial transcriptomics data into actionable scientific insights.

STAgent advances spatial transcriptomics analysis through its integrated suite of capabilities. Its autonomous code generation enables dynamic creation and execution of analytical scripts tailored to spatial transcriptomics data without human intervention, addressing a critical technical barrier. This capability was demonstrated through autonomous workflows for cell composition analysis, spatial mapping, and neighborhood analysis of SC-pancreas grafts. Additionally, leveraging vision-language models, STAgent’s visual reasoning allows it to directly interpret spatiotemporal patterns in complex tissue architectures, identifying subtle changes typically requiring weeks of expert examination. Notably, the agent identified the progressive reorganization of α- and β-cells into mature islet-like structures with distinct mantle-core arrangements and detected the expanding mesenchymal network supporting these structures.

STAgent’s contextualized biological interpretation, achieved through real-time literature search, enhances its ability to relate spatial observations to broader scientific contexts. This capability transformed spatial data into meaningful biological insights about SC-pancreas development, connecting observed patterns to functional implications supported by relevant literature. For example, the agent autonomously linked the observed α-β cell spatial arrangements to coordinated hormone secretion in mature islets described in existing studies. Furthermore, STAgent’s context-aware gene set analysis identified biologically relevant processes driving β-cell maturation, including enhanced ion transport, secretory granule formation, and cell-matrix interactions, avoiding irrelevant pathways commonly highlighted by generic enrichment tools.

STAgent’s core autonomous research capability integrates code generation, multimodal reasoning, literature-based contextualization, and biological interpretation into a seamless workflow that transforms spatial data into actionable scientific knowledge. Analytical tasks that would typically take weeks of effort from computational biologists and domain experts are completed by STAgent in minutes, dramatically reducing the expertise barrier and shortening analysis time while preserving biological relevance. By merging computational analysis with biological context and literature knowledge, STAgent serves as a valuable tool for researchers investigating tissue development and regeneration.

Looking forward, several exciting possibilities emerge for extending STAgent’s capabilities. Integration with other omics modalities, such as spatial proteomics, metabolomics or multiomics, could provide more comprehensive insights into tissue development and function. Application to larger and more diverse datasets could enable comparative analyses across different experimental conditions, disease states, or developmental trajectories.

In conclusion, we demonstrate how the combination of cutting-edge spatial transcriptomics with autonomous AI agent-driven analysis can uncover complex developmental processes with unprecedented efficiency and depth. STAgent advances our understanding of SC-pancreas graft maturation *in vivo* and establishes a new paradigm for analyzing complex biological datasets. By transforming spatial data into actionable scientific knowledge within minutes, autonomous AI agents like STAgent have the potential to accelerate discovery in spatial biology and beyond.

## Methods

### AI agent orchestration framework of STAgent

We developed STAgent using LangChain and LangGraph AI agent frameworks to establish a robust infrastructure for autonomous spatial transcriptomics analysis. These frameworks enabled the design of complex, stateful workflows essential for the multistep nature of spatial transcriptomics analysis. We leveraged advanced multimodal LLMs including Claude-3.7-Sonnet and GPT-4o for their reasoning and visual understanding capabilities. These models were used without fine-tuning to preserve their general-purpose flexibility while applied to specialized biological tasks.

The core architecture of STAgent implements a reasoning-action cycle wherein the agent sequentially: (1) observes multimodal data, (2) performs multimodal reasoning to determine appropriate analytical approaches, (3) generates and executes code, (4) evaluates results and retrieve relevant literatures, and (5) plans subsequent steps. This iterative process, guided by conditional logic and feedback loops, allows the agent to refine analyses adaptively based on intermediate findings. To ensure coherence across long-running analyses, we incorporated state persistence mechanisms that maintain analytical context through multiple reasoning steps.

### Multimodal context memory system of STAgent

We developed a specialized multimodal context memory system to enhance STAgent’s ability to maintain analytical continuity across reasoning steps. This memory system manages various forms of information, including text descriptions of analytical objectives and progress; intermediate results, such as quantitative outputs and observations; generated visualizations, including spatial maps and plots; and conversation history, preserving interaction sequences for contextual reasoning. This memory system allows STAgent to reference and reason about previously generated spatial representations and analytical outputs, supporting coherent interpretation over iterative analysis processes. Maintaining awareness of evolving analytical visualizations is particularly critical for spatial transcriptomics, where changes in tissue architecture and molecular expression often emerge gradually over multiple analysis steps.

### Code execution tool of STAgent

STAgent employs a CodeAct-inspired framework^13^ to consolidate analytical actions into a unified, flexible code action space. This approach facilitates dynamic code generation and execution, allowing the system to adapt seamlessly to various spatial transcriptomics analysis scenarios and user inputs. We implemented a Python REPL tool for executing generated code, enabling real-time interaction with data and results.

### Code knowledge retrieval system of STAgent

To enhance the general coding capabilities of multimodal LLMs with specialized spatial transcriptomics analysis techniques, we implemented a code retrieval-augmented generation (RAG) system. This system implements external code repositories from Github, processed through a segmentation pipeline that divides them into contextually meaningful chunks while preserving function definitions and related code blocks.

The segmented code is embedded using OpenAI’s embedding models and stored within a Chroma vector database, enabling efficient retrieval based on semantic similarity. During analysis, STAgent formulates queries derived from the current context and retrieves relevant code examples from this knowledge base. The retrieved code serves as a foundation that the multimodal LLM can adapt and extend to meet specific analytical requirements, enabling sophisticated spatial analyses beyond the standard capabilities of its underlying LLMs.

### Literature integration system of STAgent

We developed a real-time literature integration system to connect STAgent’s analytical capabilities with the broader scientific knowledge base. This system employs Google Scholar and SerpAPI to conduct targeted searches based on the current analytical context, retrieving relevant scientific literature to inform spatial transcriptomics interpretation.

The retrieved papers are processed to extract key information pertaining to biological mechanisms, developmental processes, and tissue architecture relevant to the analysis. This information is incorporated into STAgent’s multimodal reasoning process, allowing it to contextualize spatial patterns within established biological frameworks. The literature integration system also supports citation of relevant studies, enhancing the credibility and scientific rigor of the generated interpretations.

### Frontend of STAgent

We developed STAgent’s user interface using Streamlit, providing an intuitive, accessible platform that requires no computational expertise. The interface features an interactive chat system supporting text, voice commands, and image uploads, allowing users to interact with the agent through their preferred modality. STAgent responds with rich multimodal outputs including detailed text explanations, offering clear, contextualized insights based on agent analysis; executable code snippets, allowing users to inspect or further adapt computational workflows; interactive visualizations, facilitating the exploration of spatial data through dynamically generated figures; and comprehensive research reports, structured similarly to scientific publications. A customized sidebar enables users to save, load, and summarize research sessions, ensuring continuity across multiple interactions. Voice interaction capabilities, implemented via speech recognition and text-to-speech enhance accessibility for hands-free operation. Image uploads allow researchers to provide supplementary data directly to the agent.

State management is handled by Streamlit’s session state mechanism, ensuring analytical context persists across complex, iterative analyses. The frontend communicates seamlessly with the LangGraph-based backend via an API layer that efficiently transmits multimodal data.

Visualization components are dynamically generated through Python code created by the agent, rendering complex spatial data into accessible formats via natural language requests with minimal programming effort.

### Stem-cell derived pancreas islet differentiation

Human pluripotent stem cells (HUES8, NIH hPSC registry #09-0021) were seeded at a density of 0.6 × 10⁶ cells/mL in mTeSR 1 medium (StemCell Technologies), supplemented with 10 μM ROCK inhibitor Y27632 (DNSK International), and cultured in a 30 mL suspension bioreactor (ReproCELL) to initiate directed differentiation toward islet organoids as previously described^25,26,48^. At 24 hours post-seeding, a 15 mL half-medium exchange with fresh mTeSR 1 was performed, followed by a full 30 mL media replacement at 48 hours. At 72 hours, differentiation was initiated using a stepwise protocol. During Stage 1 (consisting of three days in S1 medium), cells were treated with 100 ng/mL Activin A and 14 μg/mL CHIR99201 on Day 1, and 100 ng/mL Activin A on Day 2, with no media change on Day 3. During Stage 2 (consisting of two days in S2 medium), cells were treated with 50 ng/mL KGF on Day 1, followed by no media change on Day 2. During Stage 3 (consisting of two days in S3 medium), cells were treated with 50 ng/mL KGF, 0.25 μM Sant1, 2 μM retinoic acid (RA), 500 nM PDBU, 10 μM ROCK inhibitor, and 200 nM LDN193189 on Day 1; LDN193189 was omitted on Day 2, while other factors were maintained. During Stage 4 (consisting of five additional days in S3 medium), cells were treated with 50 ng/mL KGF, 0.25 μM Sant1, 0.1 μM RA, 10 μM ROCK inhibitor, and 5 ng/mL Activin A, with media changes every other day (for 3 changes total). During Stage 5 (consisting of seven days in BE5 medium), cells were treated with 0.25 nM Sant1, 20 ng/mL β-cellulin, 1 μM XXi, 10 μM Alk5i II, 1 μM triiodothyronine (T3), and 0.1 μM RA from Days 1 to 4, followed by medium lacking Sant1 and containing 0.025 μM RA from Days 5 to 7; media was refreshed every other day throughout (for 4 changes total). Stage 6 consisted of extended culture in S3 medium with feeding every other day.

### Pancreas graft transplantation on mice

Human pluripotent stem cell-derived pancreatic cells were transplanted beneath the kidney capsule of immunodeficient SCID-Beige mice (Jackson Laboratory). Animals were housed under 12-hour light/dark cycles and provided standard rodent chow. Grafts were harvested at 4-, 16-, and 20-weeks post-transplantation to capture critical stages of *in vivo* differentiation and maturation. All animal procedures followed Institutional Animal Care and Use Committee (IACUC) guidelines, with transplantations performed as described by Pagliuca et al. (2014). Stem cell-derived islet organoids were harvested, resuspended in RPMI-1640 medium (Life Technologies; 11875-093), and maintained on ice before catheter implantation (5 × 10⁷ cells per mouse).

### *In situ* sequencing on pancreas grafts

Kidneys bearing pancreatic grafts were collected at 4-, 16-, and 20-weeks post-transplantation, embedded in optimal cutting temperature (OCT) compound, cryo-sectioned (20 µm), and mounted onto glass-bottom 12-well plates (P12-1.5H-N, Cellvis). Tissue sections were fixed in 4% paraformaldehyde (Electron Microscopy Sciences, 15710-S) in PBS for 15 min at room temperature, permeabilized using pre-chilled methanol (Sigma-Aldrich, 34860-1L-R) at −20°C for 1 hour, and rehydrated with PBSTR/glycine/yeast tRNA solution (PBS, 0.1% Tween-20, 0.1 U/µL SUPERase-In, 100 mM glycine, and 0.1 mg/mL yeast tRNA) at room temperature for 15 min, followed by two 5 min washes with PBSTR. Rehydrated sections were incubated in 30% formamide in 2× SSC at 30 °C for 3 hours, and in hybridization buffer (2× SSC, 1% Tween-20, 0.2 U/µL SUPERase-In, 10% formamide, 20 mM ribonucleoside vanadyl complexes, 0.1mg/mL yeast tRNA) containing SNAIL probes (5 nM per oligo) targeting a dual-species gene panel for human pancreas and mouse kidney at 40°C overnight with gentle shaking. Post-hybridization, tissues samples were twice washed in PBSTR at 37°C for 20 min, followed by a 20 min wash with high salt buffer (PBSTR, 4× SSC). Subsequently, samples were incubated overnight with a T4 DNA ligase mixture (0.1 U/µL T4 DNA ligase (Thermo Scientific, EL0011), 1× T4 ligase buffer, 0.5 mg/mL BSA (New England Biolabs, B9000S), 0.2 U/µL of SUPERase-In) at room temperature, followed by 3 washes with PBSTR, each for 5 min. Samples were then incubated with rolling-circle amplification mixture (0.2 U/µL Phi29 DNA polymerase (Thermo Scientific, EP0094), 1× Phi29 reaction buffer, 250 µM dNTP mixture (New England Biolabs, N0447S), 0.5 mg/mL BSA, 0.2 U/µL of SUPERase-In and 20 µM 5-(3-aminoallyl)-dUTP (Invitrogen, AM8439)) for 30 min at 4 °C to reach equilibrium, then for 2 hours at 30 °C to amplify. After amplification, samples were rinsed twice in PBST (PBS with 0.1% Tween-20), treated with 25 mM NHS-acrylic acid for 1 hour, then incubated in a monomer buffer (4% acrylamide, 0.2% bis-acrylamide, 0.2% TEMED, 2× SSC) for 30 min. To polymerize, a monomer buffer containing 4% acrylamide, 0.2% bis-acrylamide, 0.2% TEMED, and 0.2% ammonium persulfate was added, followed immediately by a Gel Slick-coated coverslip, after which samples were incubated for 1 hour at room temperature under nitrogen. Finally, tissue-gel hybrids were digested overnight with Proteinase K in 2× SSC with 1% sodium dodecyl sulfate (SDS) and washed thoroughly in PBST prior to sequencing.

### Sample imaging with confocal microscope

In preparation for sequencing, samples were washed twice with stripping solution (0.1% Tween-20 and 10% formamide in water) for 10 min, and five times with PBS for 5 min. Samples were then treated with 0.25 U/µL Antarctic Phosphatase (New England Biolabs, M0289L) in 1× Antarctic Phosphatase buffer containing 0.5 mg/mL BSA at 37°C for 1 hour. Sequencing by ligation was performed using a mixture containing 0.2 U/µL T4 DNA ligase, 1× T4 ligase buffer, 0.5 mg/mL BSA, 10 µM reading probe, and 5 µM fluorescence decoding probes, incubated overnight at room temperature. Following ligation, samples were washed three times with imaging buffer (10% formamide, 2× SSC in water) for 10 min.

Imaging was conducted using a Leica TCS SP8 confocal microscope equipped with a 405 nm diode laser, white-light laser, and a 40 × oil immersion objective at a voxel size of 142 nm × 142 nm × 396 nm. Images were acquired sequentially for multiple fluorescence channels corresponding to the ligation reactions.

### Image processing and cell segmentation

Raw fluorescence images were processed using a custom computational pipeline integrating MATLAB and Python. Images underwent multidimensional histogram matching, followed by top-hat filtering. Global alignment was performed using 3D fast Fourier transform-based cross-correlation. Subsequently, local alignment was achieved via non-rigid registration (using MATLAB’s imregdemons function), with the initial imaging round serving as the reference. Amplicons were detected by identifying local intensity peaks around dot centroids across four imaging channels. Gene identities were assigned by matching detected amplicon barcode sequences. Cell segmentation was accomplished using ClusterMap.

### Cell type annotation and clustering analysis

Single-cell spatial transcriptomics data from pancreas graft and kidney tissue samples were processed with the Scanpy package in Python. Data normalization was performed using sc.pp.normalize_total(), followed by log transformation with sc.pp.log1p() and scaling to unit variance and zero mean with sc.pp.scale(). Principal Component Analysis (PCA) was conducted using sc.tl.pca(). A neighborhood graph was then constructedwith sc.pp.neighbors(). Dimensionality reduction and visualization were achieved through Uniform Manifold Approximation and Projection (UMAP) using sc.tl.umap(). Unsupervised clustering was performed using the Leiden algorithm with sc.tl.leiden(), and cell types were annotated based on the expression profiles of known marker genes, including human endocrine (INS, GCG, SST, TPH1), human exocrine (CPA2, CTRB2), human vascular (COL3A1, COL1A1, SPARC, VIM), mouse ureteric epithelium (HSD11B2, AQP2), mouse endothelial/vascular (CDH5, KDR), and mouse nephron (SPP2, LRP2).

### Spatial analysis and domain identification

To identify spatial domains in pancreas graft samples across multiple timepoints, we utilized the STAligner algorithm for integration of spatial transcriptomics data. Spatial networks were constructed individually for each tissue section using STAligner.Cal_Spatial_Net(). Section-specific AnnData objects were concatenated into a single dataset using ad.concat(), and adjacency matrices were combined into a block diagonal matrix using scipy.sparse.block_diag(). We trained STAligner using a graph attention network architecture initialized with STAGATE embeddings to align all tissue sections. Subsequently, unsupervised Louvain clustering was applied to the aligned embeddings, and spatial domains were manually annotated based on biological relevance.

### Gene imputation with Tangram

We performed whole-transcriptome gene imputation using Tangram to map scRNA-seq (GSE151117) data onto spatial coordinates. Prior to mapping, datasets were harmonized for compatibility by assigning unique cell identifiers in the scRNA-seq data and retaining only human cells in spatial datasets. Marker genes were identified through differential expression analysis sc.tl.rank_genes_groups(), and datasets were normalized using sc.pp.normalize_total(). Tangram’s preprocessing function tg.pp_adatas() identified genes for model training. Cell mapping was executed with tg.map_cells_to_space(), followed by projection of complete gene expression profiles onto spatial data via tg.project_genes().

### Cell-cell communication profiling with CellChat

Cell-cell communication within pancreas grafts at various timepoints were profiled using CellChat applied toTangram-imputed spatial transcriptomics data. We utilized the human-specific CellChatDB, excluding non-protein signaling pathways. Data were subset to signaling-related genes with subsetData(), and stochasticity in expression was addressed using the smoothData() function based on the human protein-protein interaction (PPI) network. Cell-cell communication networks were inferred through a sequential pipeline: identifying overexpressed genes and signaling interactions, computing communication probabilities using the trimean method, and filtering to retain interactions involving cell types represented by at least five cells. Pathway-level analyses were completed using computeCommunProbPathway() and aggregateNet(). Comparative analyses across timepoints were performed by generating pairwise combinations of CellChat objects, visualized through comparative bar plots, differential network plots, and heatmaps. This approach revealed conserved and timepoint-specific intercellular signaling patterns relevant to pancreas graft development.

## Code availability

Software code for this study will be made publicly available and maintained at the time of publication at http://github.com/LiuLab-Bioelectronics-Harvard/STAgent.

## Acknowledgments

We acknowledge support from NIH/NIDDK 1DP1DK130673 (J.L. and J.RA.-D), NSF ECCS-2038603 (J.L.), NIH/NLM 5R01LM014465 (J.L. and J.D.). Edward Scolnick Professorship (X.W.), Packard Fellowship for Science and Engineering (X.W.), and NIH New Innovator Award (X.W.).

## Author contributions

Z.L., W.W., J.RA.-D. and J.L. conceived the idea. Z.L. and W.W. developed the STAgent method. J.RA.-

1. D. prepared SC-pancreas graft mice. Z.L., Q.L. and X.S. performed the STARmap experiment. Z.L. and

W.W. prepared figures and drafted the manuscript. All authors contributed critical discussions and input on the figures and results. J.RA.-D. and J.L. supervised the study.

## Competing interest statement

J.L. is cofounder of Axoft, Inc., and X.W. is a scientific cofounder and equity holder of Stellaromics, Inc.

---

**Extended Data Fig. 1:**
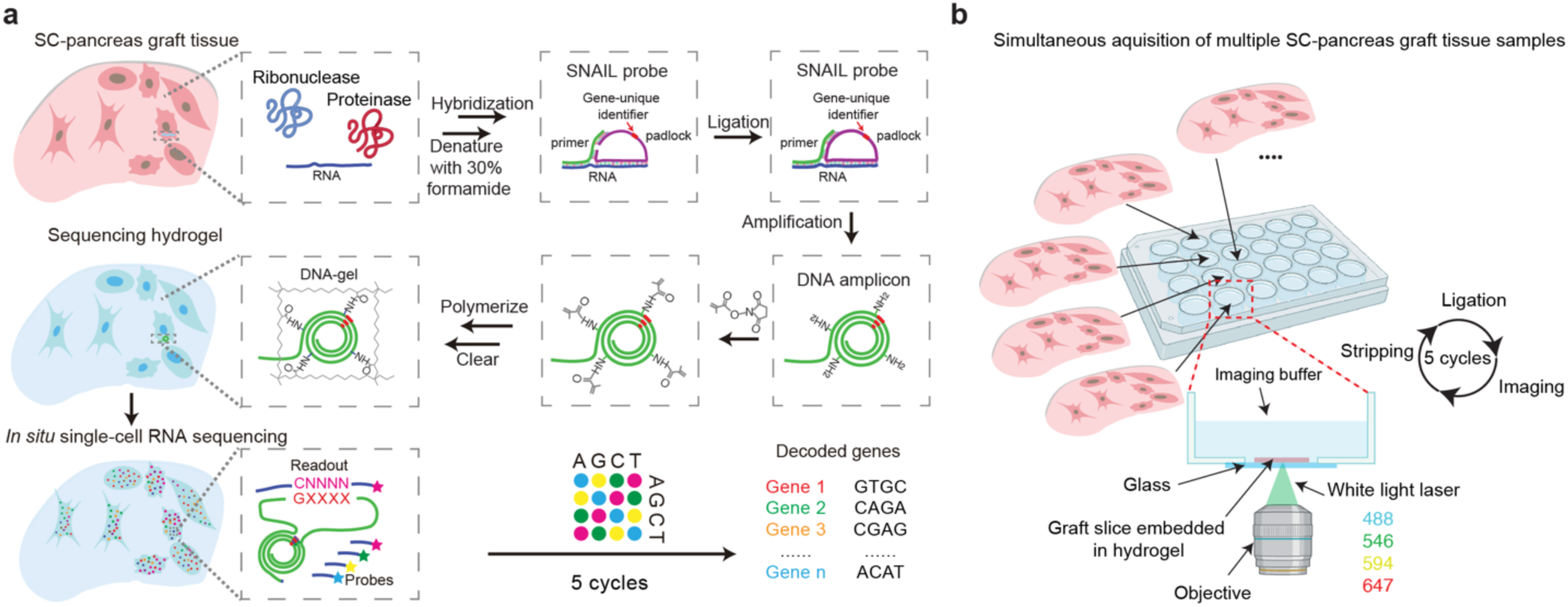
Optimized spatial transcriptomics methodology in SC-pancreas grafts. **(a)** Modified STARmap protocol optimized for pancreatic tissue, involving extended mRNA denaturation (>3 hours) and increased formamide concentration (30%) during hybridization. This is followed by DNA amplicon generation, sequencing, and gene decoding across five imaging cycles. (**b**) Experimental setup enabling simultaneous acquisition of multiple SC-pancreas graft tissue samples.

**Extended Data Fig. 2:**
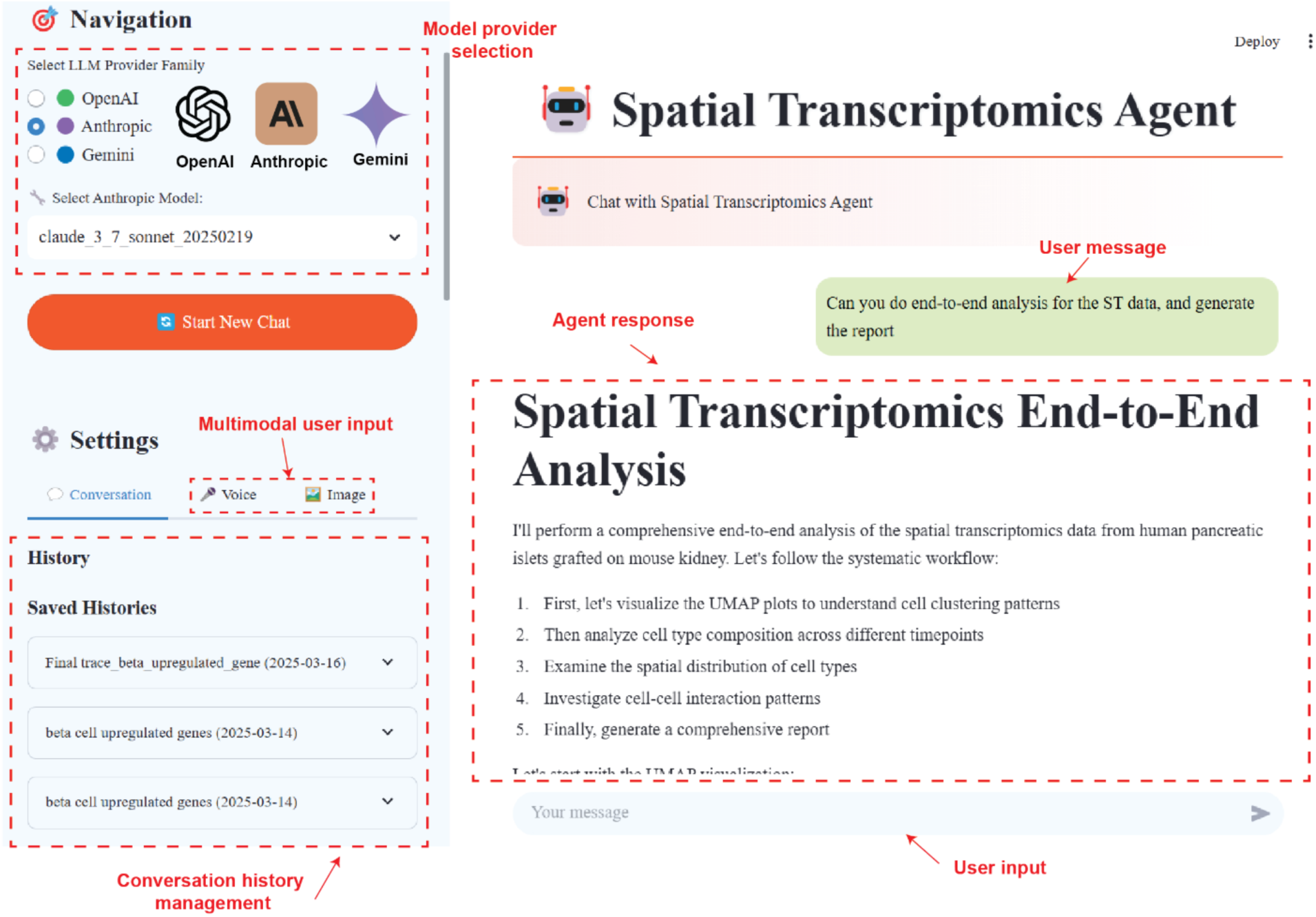
STAgent user interface for spatial transcriptomics analysis. STAgent features a Streamlit-based frontend designed for intuitive spatial transcriptomics analysis through conversational interactions. Interface components include: model provider selection (top left), allowing users to choose between different LLM providers (OpenAI, Anthropic, Gemini) and specific models; multimodal user input**s** (middle left), supporting text, voice, and image uploads; conversation history management (bottom left), enabling users to save and revisit previous analyses; and the interaction panel (right), where users can submit natural language queries and receive detailed responses. This interface enables researchers without computational expertise to conduct advanced spatial transcriptomics analyses effortlessly.

**Extended Data Fig. 3:**
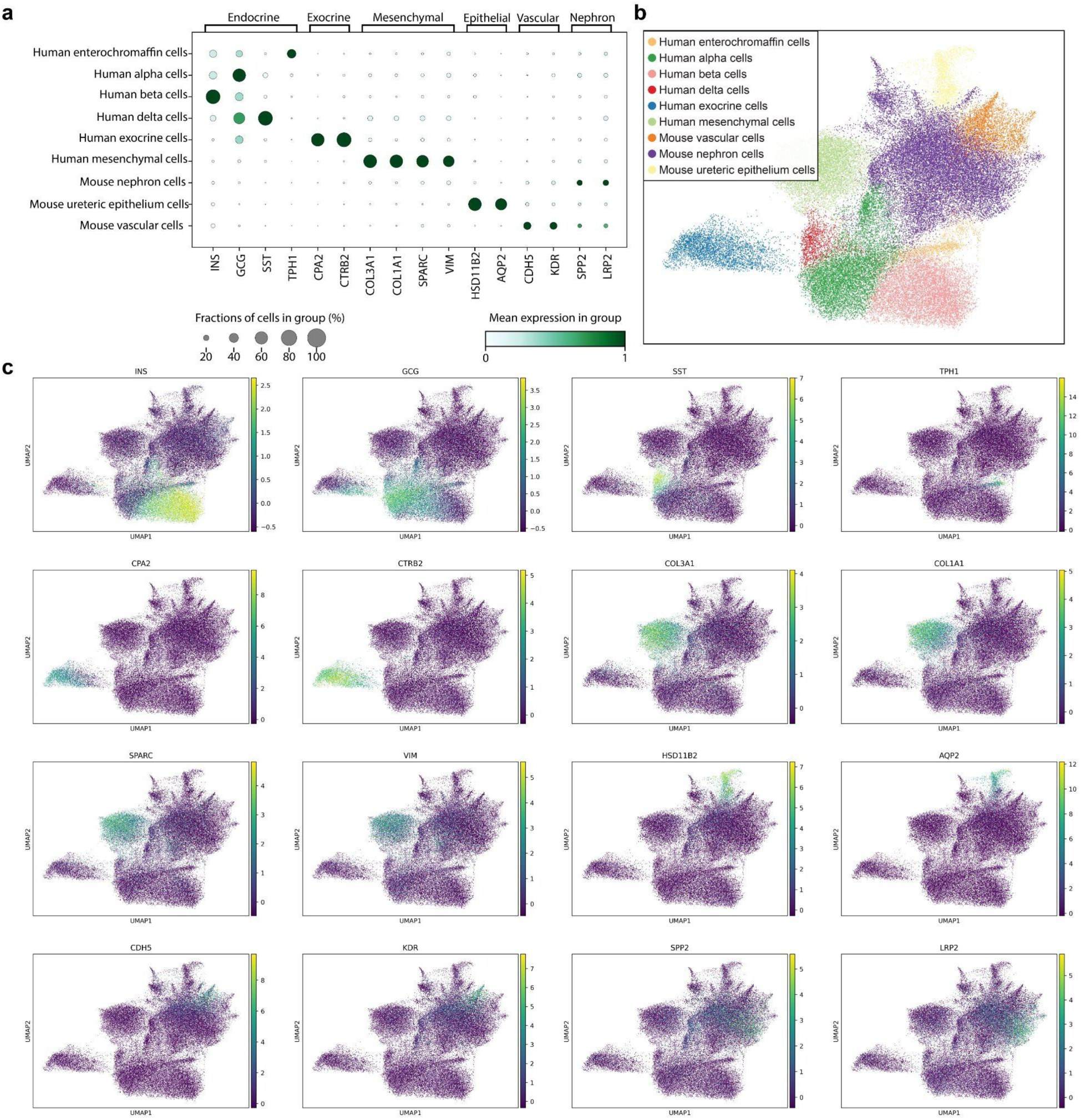
Cell type annotation and marker gene expression profiling. (**a**) Dot plot illustrating key marker gene expression profiles across annotated cell types. Dot size represents the fraction of expressing cells in the graft, while color intensity indicates mean gene expression levels. (**b**) UMAP embedding of all cells colored by assigned cell type, highlighting distinct cell population clusters (**c**) UMAP feature plots for selected marker genes, showing their spatial expression patterns within identified clusters.

**Extended Data Fig. 4:**
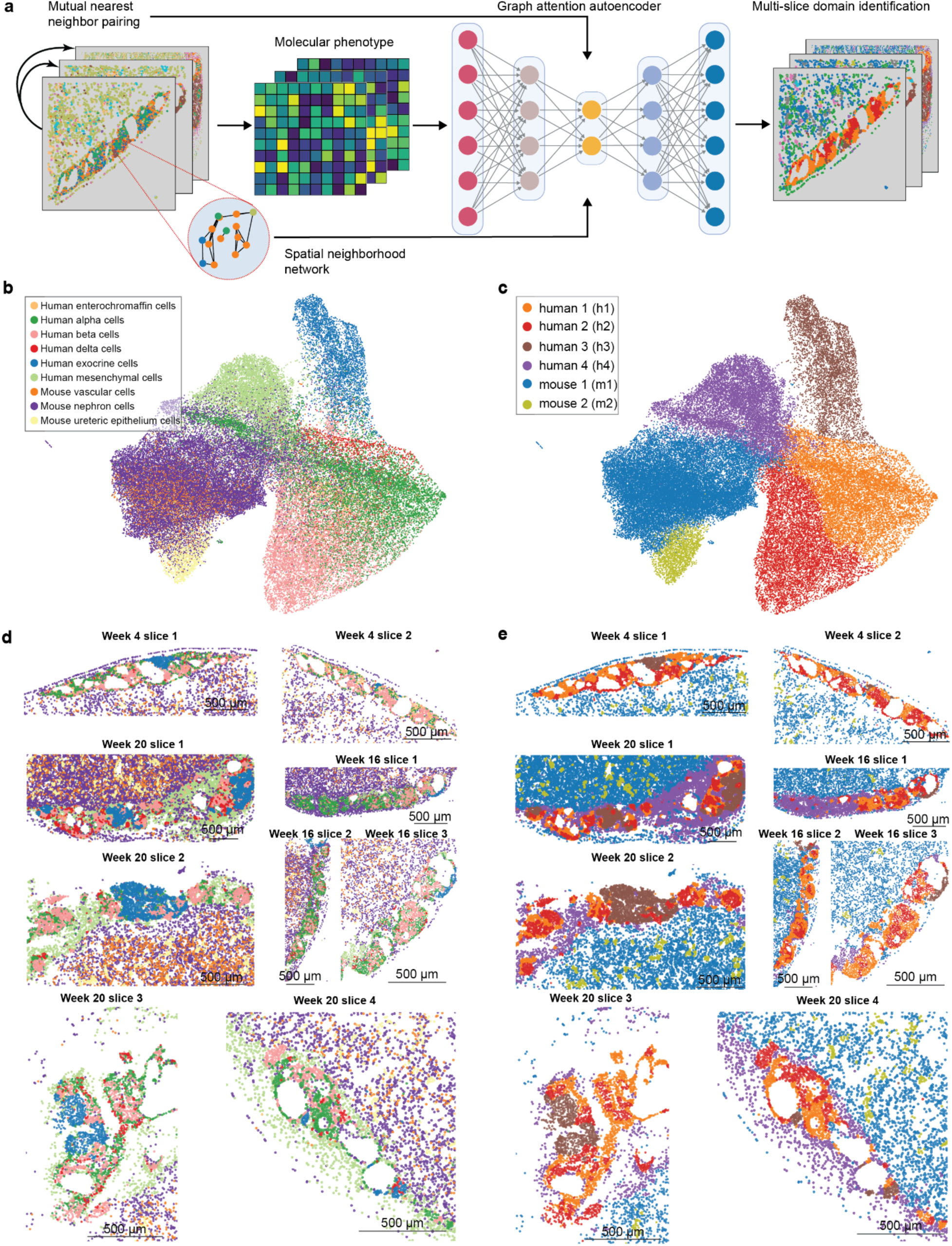
Multi-slice domain identification. **(a)** Pipeline schematic for identifying spatial domains across multiple tissue slices. A graph attention autoencoder integrates molecular features and spatial neighborhood networks, facilitating automated domain segmentation. (**b-c**) STAligner embeddings for each tissue slice colored either by cell type (**b**) or by assigned spatial domain (**c**), demonstrating consistent domain identification across slices. (**d**) Composite spatial cell type map depicting the distribution of each identified cell type across all analyzed tissue sections. (**e**) Corresponding spatial domain map highlighting the locations of each domain in each tissue slice.

**Extended Data Fig. 5:**
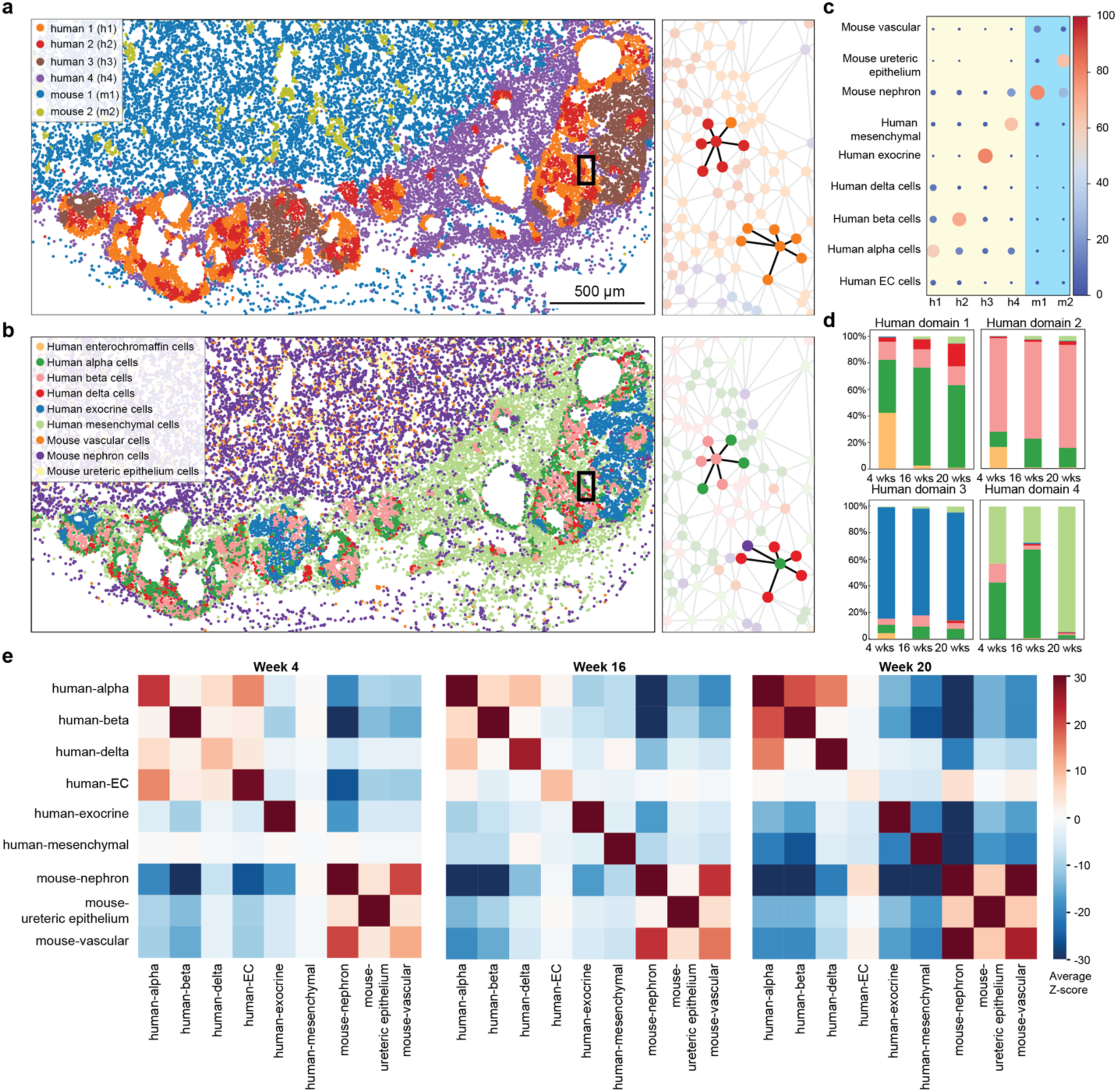
Sptial domain identification and cell-cell neighborhood enrichment analysis. (a-b) Representative spatial maps showing identified spatial domains (**a**) and corresponding cell type distributions (**b**) in pancreas grafts. Network diagrams (right) illustrate cell adjacency and highlight local tissue architecture. **(c)** Dot chart displaying domain-specific cell-type composition, revealing relative abundances of major cell populations within each domain. **(d)** Bar plots illustrating changes in the proportion of each spatial domain at 4-, 16-, and 20-weeks post-transplantation. **(e)** Heatmaps depicting spatial neighborhood enrichment at 4-, 16-, and 20-weeks post-transplantation.

**Extended Data Fig. 6:**
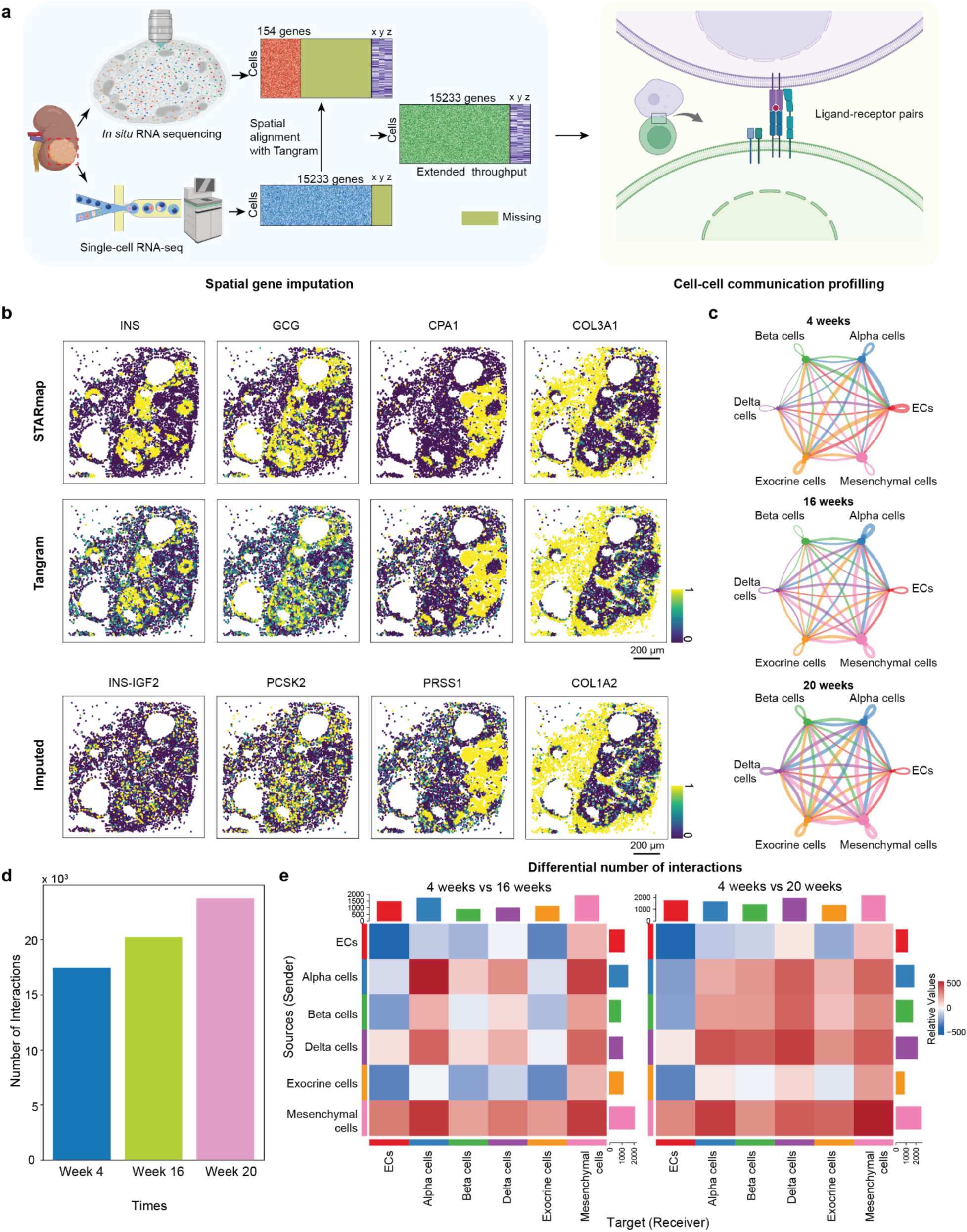
Whole-transcriptome imputation of spatial transcriptomics data and cell-cell communication analysis. **(a)** Workflow schematic for Tangram-based imputation. Published scRNA-seq data are integrated with *in situ* transcriptomics to infer whole-transcriptome gene expression. **(b)** Spatial expression maps of selected marker genes (e.g., INS, GCG, CPA1, COL3A1), demonstrating consistent localization for both observed and newly imputed genes (INS-IGF2, PCSK2, PRSS1, and COL1A2). **(c)** Flow diagram summarizing inferred cell-cell communication networks at 4-, 16-, and 20-weeks post-transplantation, highlighting interaction numbers. **(d)** Bar plot depicting the total number of inferred interactions at each analyzed time point. **(e)** Heatmap comparing the differential number of interactions between cell types at 4-vs. 16-weeks and 4- vs. 20-weeks post-transplantation.

